# Dimerization activates the Inversin complex in *C. elegans*

**DOI:** 10.1101/2024.05.17.594761

**Authors:** Erika Beyrent, Derek T. Wei, Gwendolyn M. Beacham, Sangwoo Park, Jian Zheng, Matthew J. Paszek, Gunther Hollopeter

**Affiliations:** Department of Molecular Medicine, Cornell University, Ithaca, NY; Field of Biochemistry, Molecular, and Cell Biology, Cornell University, Ithaca, NY; Field of Biophysics, Cornell University, Ithaca, NY; Robert Frederick Smith School of Chemical and Biomolecular Engineering, Cornell University, Ithaca, NY; Nancy E. and Peter C. Meinig School of Biomedical Engineering, Cornell University, Ithaca, NY

**Author notes:** Correspondence and requests for materials should be addressed to G.H. these authors contributed equally (Derek T. Wei, Gwendolyn M. Beacham). Department of Medicine, Section of Hematology and Medical Oncology, Boston University Chobanian & Avedisian School of Medicine and Boston Medical Center, Boston, MA. Molecular Biology Program, Memorial Sloan Kettering Cancer Center; and Programs in Biochemistry, Cell, and Molecular Biology, Weill Cornell Graduate School of Medical Sciences, New York, NY. The authors declare no competing interests.

## Abstract

Genetic, colocalization, and biochemical studies suggest that the ankyrin repeat-containing proteins Inversin (INVS) and ANKS6 function with the NEK8 kinase to control tissue patterning and maintain organ physiology. It is unknown whether these three proteins assemble into a static “Inversin complex” or one that adopts multiple bioactive forms. Through characterization of hyperactive alleles in *C. elegans*, we discovered that the Inversin complex is activated by dimerization. Genome engineering of an RFP tag onto the nematode homologs of INVS (MLT-4) and NEK8 (NEKL-2) induced a gain-of-function, cyst-like phenotype that was suppressed by monomerization of the fluorescent tag. Stimulated dimerization of MLT-4 or NEKL-2 using optogenetics was sufficient to recapitulate the phenotype of a constitutively active Inversin complex. Further, dimerization of NEKL-2 bypassed a lethal MLT-4 mutant, demonstrating that the dimeric form is required for function. We propose that dynamic switching between at least two functionally distinct states—an active dimer and an inactive monomer—gates the output of the Inversin complex.

## Introduction

Genetic data suggest that the ankyrin repeat-containing protein Inversin (INVS) functions with at least two other proteins, the never-in-mitosis A-related kinase 8 (NEK8) and the ankyrin repeat and sterile alpha motif-containing protein 6 (ANKS6). Loss of any one of the three proteins results in similar pathologies in humans and vertebrate models. INVS was initially identified at the genetic locus disrupted in mice that exhibit neonatal mortality with reversal of left-right asymmetry (*situs inversus*) (Yokoyama et al. 1993; Mochizuki et al. 1998; Morgan et al. 1998). INVS was later found to also be mutated in human patients presenting with infantile nephronophthisis – a disease characterized by *situs inversus* and end stage renal failure, with cysts and fibrosis affecting multiple organs (Otto et al. 2003; Zhong et al. 2022). Two additional nephronophthisis loci have been identified as having mutations in NEK8 and ANKS6; vertebrate models confirm that loss of either of these proteins phenocopies loss of INVS (Frank et al. 2013; Otto et al. 2008; Hassan et al. 2020; Hoff et al. 2013; Taskiran et al. 2014; Kulkarni et al. 2020; Zhong et al. 2022). In *C. elegans*, loss-of-function mutations in the INVS homolog, MLT-4, result in a lethal molting phenotype, where worms fail to shed their collagen-based cuticles (Lažetić and Fay 2017). Mirroring the genetic data from vertebrates, loss of NEKL-2 (NEK8) and MLT-2 (ANKS6) exhibit the same molting-defective phenotype (Yochem et al. 2015; Lažetić and Fay 2017). Thus, these three proteins appear to work together across multiple metazoan species.

Although loss-of-function experiments have been crucial in revealing the biological importance of INVS, NEK8, and ANKS6, it has proven difficult to determine the precise mechanisms by which these proteins are acting to control tissue patterning and maintain organ physiology. In *C. elegans*, all three proteins have been shown to function in endocytic trafficking to regulate molting (Joseph et al. 2020, 2023). In vertebrates, a recent study proposed that INVS signals originating from the cilia counteract a cyst-activating pathway (Li et al. 2023). Previous studies have implicated INVS in non-canonical *Wnt* signaling (Simons et al. 2005; Jenny et al. 2005; Feiguin et al. 2001), while NEK8 and ANKS6 have been proposed to regulate Hippo signaling effectors (Grampa et al. 2016; Airik et al. 2020; Schwarz et al. 2022). These mechanisms have been difficult to clarify, in part, due to a lack of gain-of-function analyses that could discriminate between direct actions of these proteins and indirect pathologies associated with loss-of-function lethality.

Cellular and biochemical data support the model that INVS, NEK8, and ANKS6 constitute an ‘Inversin complex’; however, we lack fundamental knowledge about how the components might assemble into a functionally active state. Evidence from both vertebrates and invertebrates demonstrates the interdependence of the proteins for their localization. INVS appears to function as an anchor for the other two members in cilia, where the complex is observed to form a fibrillar structure by super-resolution microscopy (Bennett et al. 2020; Shiba et al. 2010). Similar to the vertebrate proteins, the *C. elegans* homologs colocalize, albeit at epithelial junctions rather than cilia (Lažetić and Fay 2017). While these localization patterns might not represent direct interactions, biochemical analyses have revealed the potential for the individual components to interact. For example, ANKS6 is reported to bind and stimulate the kinase activity of NEK8 (Nakajima et al. 2018; Czarnecki et al. 2015). It remains unclear whether the complex is a single static arrangement of the three proteins or undergoes dynamic switching between multiple states to specify its activity.

Here, we report that the Inversin complex is activated by dimerization. We identified an overt, morphological, cyst-like phenotype in *C. elegans* that signifies the complex is constitutively active. Tagging MLT-4 or NEKL-2 with a red fluorescent protein induced this phenotype, and an unbiased genetic screen for phenotype suppression yielded monomerizing mutations in the fluorescent tag. We further demonstrate that optogenetic-induced dimerization is sufficient to generate the gain-of-function phenotype. Dimerization of the Inversin complex also appears to be required for function, as we can rescue a lethal MLT-4 mutant with dimerized NEKL-2. We propose that Inversin complex activity depends on switching between inactive monomeric and active dimeric states.

## Results

### Jowls phenotype reports a constitutively active Inversin complex

We reported previously that dominant, gain-of-function alleles of *mlt-4* cause a “jowls” phenotype in which fluid-filled pockets form at the anterior ends of *C. elegans* (Beacham et al. 2022). One of these alleles was a missense mutation isolated in a forward genetic screen for jowls (MLT-4 E470K, Figure 1A and 1B). Curiously, the other allele resulted from attaching a tagRFP-T (henceforth referred to as “RFP tag”) to the C-terminus of wild-type MLT-4 (MLT-4::RFP, Figure 1A and 1B), while no phenotype resulted from tagging MLT-4 with mScarlet (Beacham et al. 2022). Although the MLT-4::RFP animals exhibit a fitness defect (Figure 1C), the phenotype is distinct from null mutants, which are lethal (Lažetić and Fay 2017), and from hypomorphic mutants, which suppress jowls (Joseph et al. 2020), suggesting that the RFP tag might instead stimulate the Inversin complex. To test this, we introduced a mutation into another member of the complex. The R12I mutation in NEK8 is associated with renal disease in humans and is hypothesized to reduce kinase activity (Hassan et al. 2020). Indeed, engineering the analogous mutation in NEKL-2 suppressed the jowls phenotype of both MLT-4 alleles (Figure 1A and 1B) and the fitness defect of MLT-4::RFP animals (Figure 1C). These results suggest that these MLT-4 alleles represent a gain-of-function that is dependent on NEKL-2.

**Figure 1:**
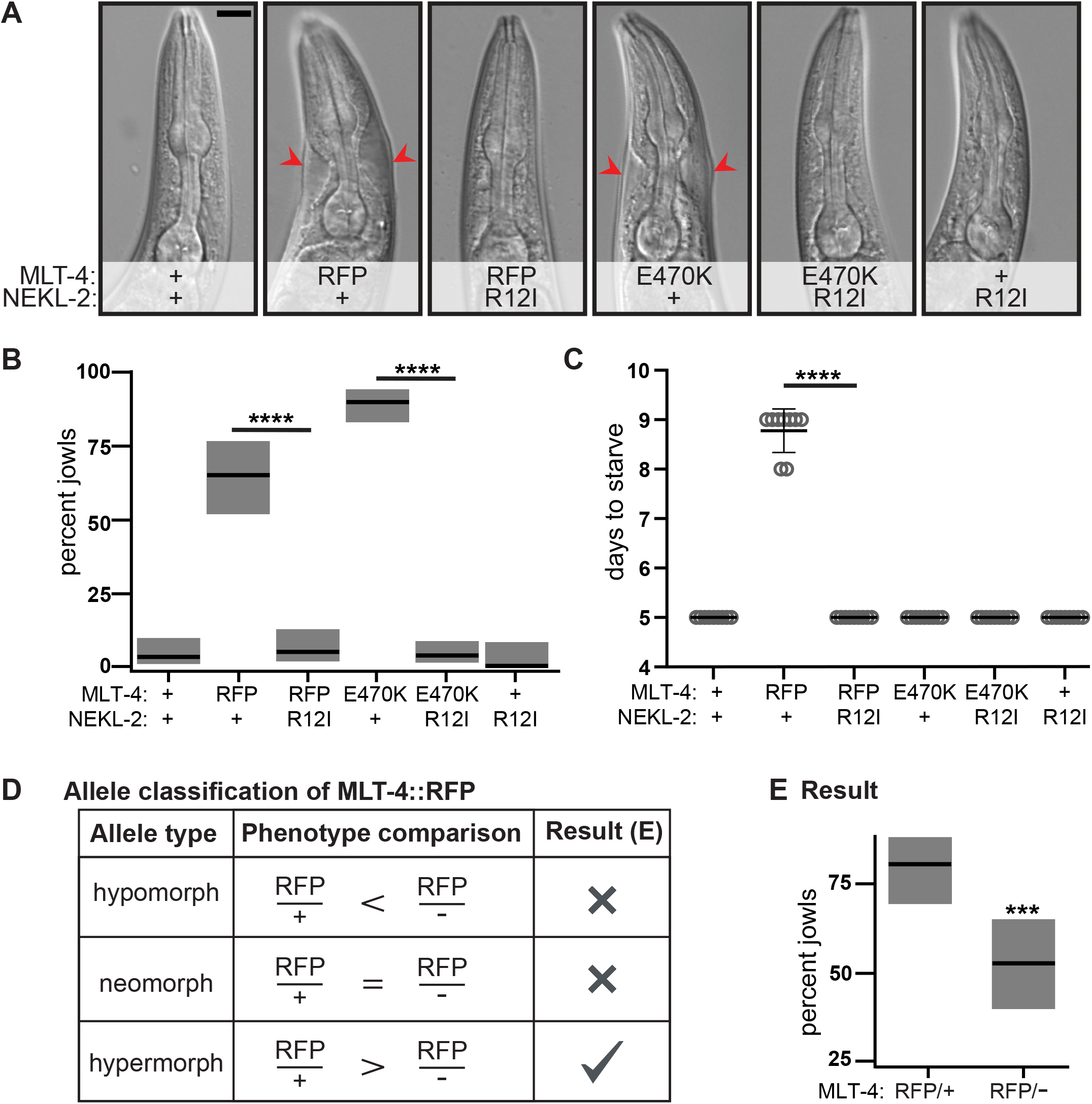
Hyperactive MLT-4 allele exhibits gain-of-function jowls phenotype. (A) Images of worm heads. Red arrows mark jowls. Scale bar = 20 μm. (B) Jowls assay. Percent of worms exhibiting jowls; gray boxes represent 95% confidence intervals. n=55-134. (C) Fitness assay. Number of days for population to expand and consume food source. Data represent mean +/-standard deviation of 10 biological replicates. (D) Predicted penetrance of jowls for each allele class. (E) Results of allele classification. Percent of worms exhibiting jowls; gray boxes represent 95% confidence intervals. + = wild-type at indicated locus, -= *mlt-4* deletion allele. n= 57-68. *** p < 0.001, **** p<0.0001, ANOVA analysis with Tukey’s post hoc test, as indicated (B,C) or compared to RFP/+ (E).

Gain-of-function alleles can result from increased protein activity (hypermorphic) or a new, unrelated activity (neomorphic). To characterize the effect of the RFP tag, we generated heterozygous strains containing one copy of MLT-4::RFP and one copy of either a wild-type MLT-4 (referred to as MLT-4::RFP/+) or a deletion allele (referred to as MLT-4::RFP/–). In this scheme (Figure 1D), hypermorphic alleles will exhibit a *reduced* phenotype in the presence of a null allele, while the phenotype of hypomorphic alleles will be *enhanced* (Muller 1932).

Neomorphic alleles act independently and will exhibit the *same* phenotype regardless of the presence of a wild-type or null allele. We found that MLT-4::RFP/+ animals had an increased penetrance of jowls as compared to MLT-4::RFP/– animals (Figure 1E). This result is consistent with MLT-4::RFP being a hypermorphic allele. To test if an RFP tag also induces a hyperactive form of the kinase NEKL-2, we generated an inducible transgene of RFP-tagged NEKL-2 (RFP::NEKL-2, Figure Supplement 1A). While expression of this transgene was lethal to larval animals, expression later in development induced jowls in adults (Figure Supplement 1B). To test whether the jowls phenotype was dependent on the kinase activity of NEKL-2, we generated a kinase-dead version (D137N; Zalli, Bayliss, and Fry 2012), and found that this mutation suppressed both the jowls phenotype and the fitness defect, as did the R12I disease allele (Figure Supplement 1B and 1C). Altogether, these results indicate that the jowls phenotype represents a biological output of a constitutively active Inversin complex.

### Disruption of the RFP dimer interface suppresses jowls

To gain insight into how the RFP tag on MLT-4 acts as a hypermorphic allele, we conducted a chemical mutagenesis screen on the MLT-4::RFP strain for animals with suppressed jowls, and isolated 17 independent amino acid changes in the RFP tag (Figure 2A). Because *mlt-4* nulls are lethal, the mutations isolated in our screen are not simply loss-of-function. Rather, we hypothesized that many of these mutations might destabilize the fluorophore and result in loss of fluorescence. Indeed, our screen selected for a mutation in the key threonine residue (T163I) that is important for photostability of tagRFP-T (Shaner et al. 2008); the crystal structure (R. Liu et al. 2016) shows this residue hydrogen bonding with the chromophore. As anticipated, our T163I mutant did not appear to be fluorescent (Figure 2A, 2D, and 2E).

**Figure 2:**
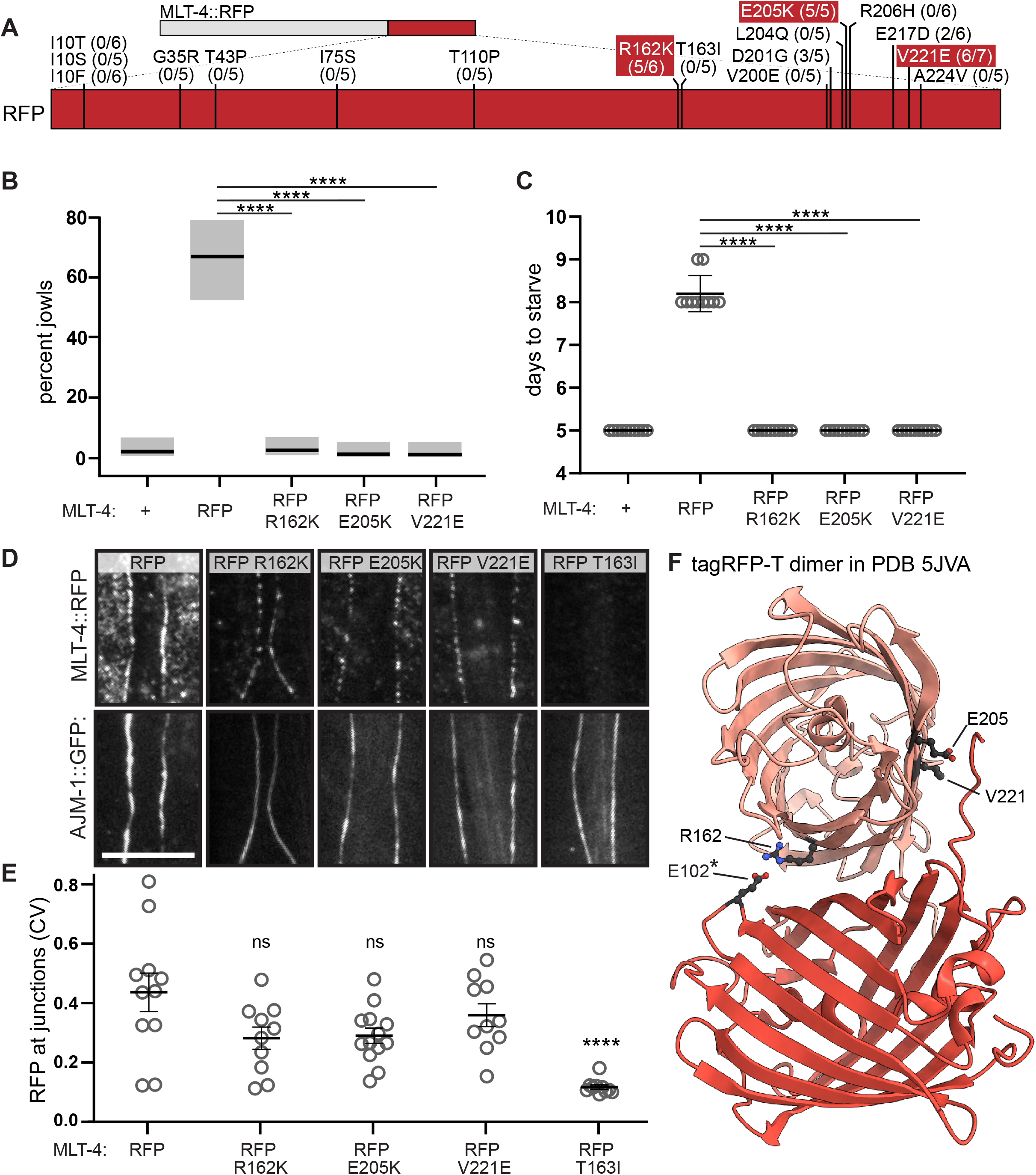
Characterization of missense mutations in RFP tag that suppress jowls. (A) Schematic of MLT-4::RFP. Fraction of worms exhibiting red fluorescence at epidermal junctions indicated for each mutant. Fluorescent mutants (red, >80%) were engineered de novo for B-E. (B) Jowls assay. Percent of worms exhibiting jowls; gray boxes represent 95% confidence intervals. n=46-138. (C) Fitness assay. Number of days for population to expand and consume food source. Data represent mean +/-standard deviation of 10 biological replicates. (D) Imaging assay for MLT-4 localization at epithelial junctions. Representative images of endogenously tagged MLT-4::RFP (top) and apical junction marker AJM-1::GFP (bottom). Scale bar = 5 μm. (E) Quantification of MLT-4::RFP pixel intensity at junctions plotted as coefficient of variance (CV). Data represent mean +/-SEM of 10-13 biological replicates. (F) Mutated residues mapped onto the structure of RFP tag (PDB 5JVA; note: residue numbers shifted -5 in PDB). * Residue predicted to form salt bridge, see Figure 3. For B and C, + = wild-type at mlt-4 locus. ns (not significant) p>0.05, **** p<0.0001, ANOVA analysis with Tukey’s post hoc test, as indicated (B,C) or compared to RFP (E).

In contrast, three of the RFP missense mutants (R162K, E205K, and V221E) retained red fluorescence (Figure 2A). To confirm that these three mutations were indeed responsible for suppression of the phenotype, we introduced them *de novo* using CRISPR. All three missense mutations suppressed both the jowls (Figure 2B) and the fitness defect (Figure 2C) of the MLT-4::RFP animals. We also confirmed that these MLT-4::RFP mutant proteins were appropriately localized by comparing their localization to that of the apical epithelial junction marker AJM-1::GFP (Z. Liu et al. 2005). Although their fluorescent signals appeared slightly reduced, all three retained a normal localization pattern (Figure 2D and 2E), suggesting these mutations might have specifically disrupted a protein-protein interaction responsible for the jowls phenotype.

When we examined the positions of these mutated residues (R162, E205, and V221) on the crystal structure of tagRFP-T (R. Liu et al. 2016), we observed that all three mapped to the dimer interface between two RFP molecules (Figure 2F). The residues analogous to E205 and V221 were previously modified to engineer monomeric forms of GFP (Zacharias et al. 2002; Costantini et al. 2012; Pédelacq et al. 2006; Scott et al. 2018). The other residue, R162, has previously been mutated to monomerize mKate (Shemiakina et al. 2012) and mCardinal (Wannier et al. 2018). To determine whether the R162K mutation isolated in our screen likewise favored MLT-4::RFP monomers, we employed single-molecule TIRF microscopy coupled with stepwise photobleaching. We lysed MLT-4::RFP worms and quantified the percentage of RFP-positive molecules within the lysate that exhibited a single photobleaching step. The R162K mutant consistently increased the monomeric fraction even as we increased the particle density (Figure 3A), consistent with an increase in RFP monomers.

**Figure 3.**
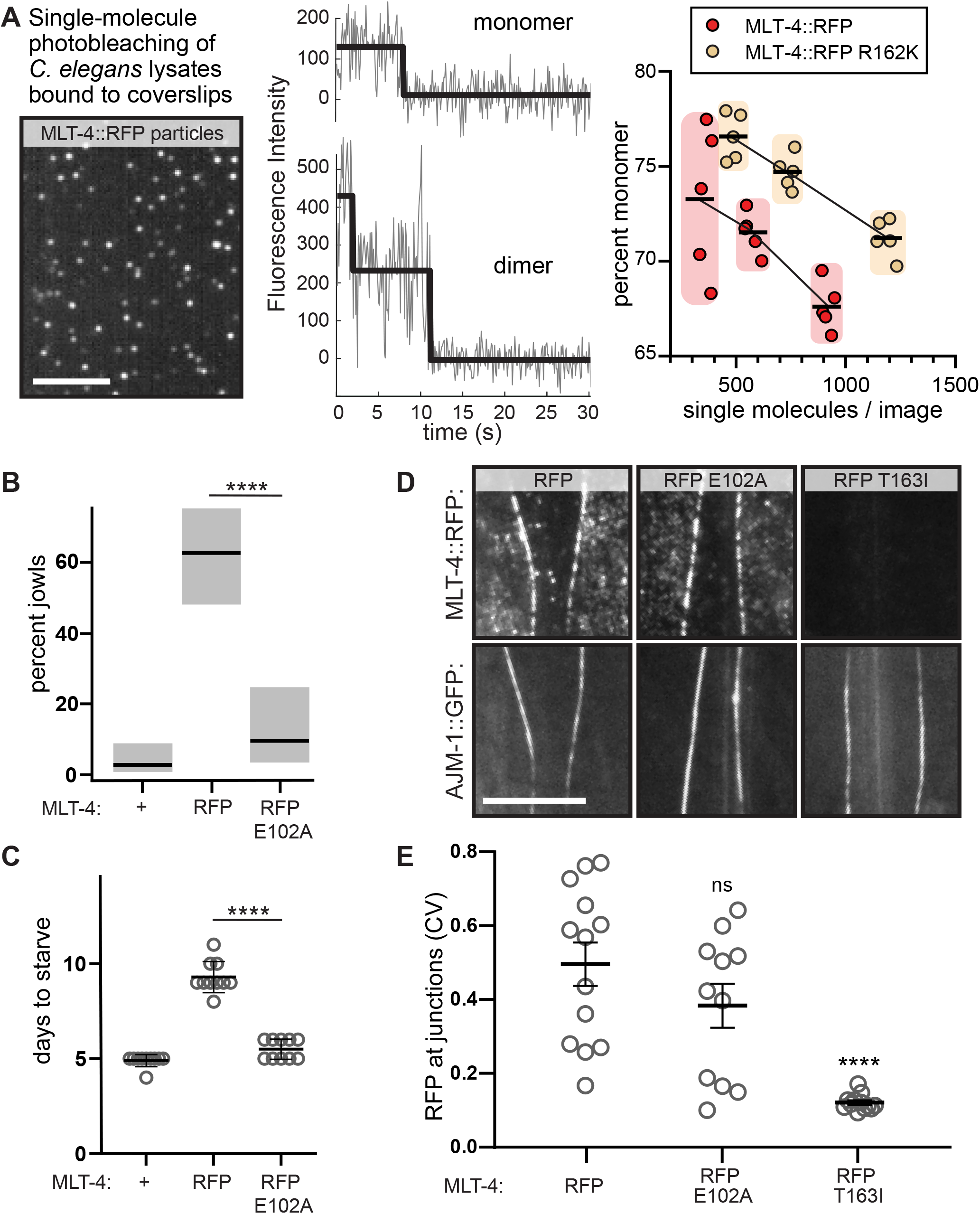
Perturbation of RFP dimerization reverses jowls phenotype. (A) Single molecule photobleaching assay. Left: Representative TIRF image of MLT-4::RFP single molecules (average intensity projection of frames 6-50). Middle: Representative fluorescence intensity traces of MLT-4::RFP monomer (top) and dimer (bottom). Right: Percent of MLT-4::RFP spots exhibiting a single bleaching step. Data collected from 5-8 images (circles) across three comparable particle densities (shaded clusters, mean indicated). (B) Jowls assay. Percent of worms exhibiting jowls; gray boxes represent 95% confidence intervals. n=35-92. (C) Fitness assay. Number of days for population to expand and consume food source. Data represent mean +/-standard deviation of 10 biological replicates. (D) Imaging assay for MLT-4 localization at epithelial junctions. Representative images of endogenously tagged MLT-4::RFP (top) and apical junction marker AJM-1::GFP (bottom). (E) Quantification of MLT-4::RFP pixel intensity at junctions plotted as coefficient of variance (CV). Data represent mean +/-SEM of 11-13 biological replicates. + = wild-type at *mlt-4* locus. ns p>0.05, **** p<0.0001, ANOVA analysis with Tukey’s post hoc test, as indicated (B,C) or compared to RFP (E). All scale bars = 5 μm.

In the crystal structure, R162 appears to bridge the dimer interface via an electrostatic interaction with residue E102 (Figure 2F). We hypothesized that mutating E102 in MLT-4::RFP animals should reduce dimerization and suppress jowls, similar to mutation of R162. Indeed, an alanine mutant (E102A) suppressed both the jowls (Figure 3B) and fitness defect (Figure 3C) of MLT-4::RFP animals, without overtly perturbing the fluorescence or localization of MLT-4::RFP (Figure 3D and 3E). Taken together, these data strongly suggest that the RFP tag forces dimerization of MLT-4 to generate a constitutively active Inversin complex.

### Optogenetic dimerization of MLT-4 and NEKL-2 causes jowls

To test the differential effects of dimerization on Inversin complex subunits, we utilized the fungal photoreceptor Vivid, which has been shown to dimerize in a light-inducible manner (Zoltowski and Crane 2008; Shrode et al. 2001). We fused Vivid to each of the Inversin complex members using CRISPR, and induced dimerization of the Vivid tag with ambient light. When we attached Vivid to the C-terminus of either MLT-4 or NEKL-2, worms exposed to light exhibited jowls as adults, while worms grown in the dark did not (Figure 4A-4C). However, we did not see a robust fitness defect associated with fusing Vivid to NEKL-2 or MLT-4 (Figure 4D), in contrast to the RFP tags. Perhaps Vivid dimerization is not as robust as RFP at overcoming potential steric barriers or regulatory mechanisms that stabilize Inversin complex monomers, or it could be that the jowls and fitness defects are due to differential outputs of the Inversin complex. Interestingly, we failed to observe a phenotype when we fused Vivid to either the N-or C-terminus of MLT-2. These results could suggest that steric hindrance prevents Vivid-dependent dimerization of MLT-2, or that MLT-2 regulates the complex through a different mechanism. Altogether, these optogenetic studies show that dimerization of either MLT-4 or NEKL-2 is sufficient to activate the complex.

**Figure 4:**
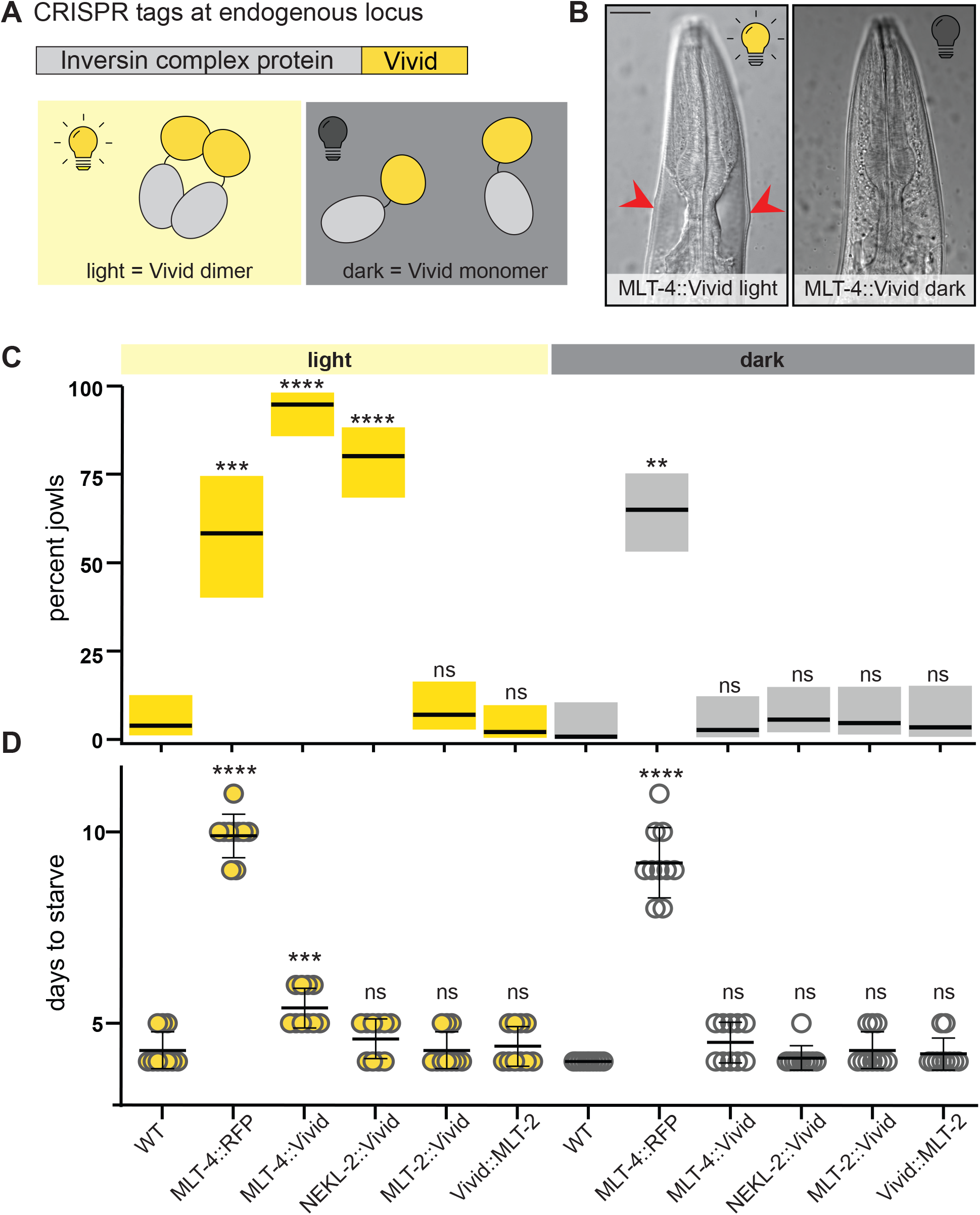
Optogenetic dimerization of MLT-4 or NEKL-2 causes jowls. (A) Diagram of light-dependent dimerization of Vivid tag fused to Inversin complex protein. (B) Images of worm heads. Red arrows indicate jowls. Scale bar = 20 μm. (C) Jowls assay. Percent of worms exhibiting jowls; gray boxes represent 95% confidence intervals. n=29-72. (D) Fitness assay. Number of days for population to expand and consume food source. Data represent mean +/-standard deviation of 10 biological replicates. WT = wild-type Bristol N2. ns p>0.05,** p<0.01, *** p < 0.001, **** p<0.0001, ANOVA analysis with Tukey’s post hoc test, compared to + within respective light/dark condition.

### Dimerization of NEKL-2 rescues a functionally inactive MLT-4 mutant

Our screen for suppressors of the MLT-4::RFP phenotype also yielded mutations in MLT-4 itself (T94P, E291A, and N442K) (Figure 5A). To determine whether any of these mutations specifically counteract dimerization, we followed the same strategy that we used to identify monomerizing mutants in the RFP tag. We engineered these MLT-4 mutations *de novo* to confirm that they suppressed MLT-4::RFP animals. While all three mutations fully suppressed the fitness defect caused by MLT-4::RFP (Figure 5B), only two (T94P and E291A) fully suppressed the jowls (Figure 5C). While the E291A mutant localized appropriately, both the N442K and T94P mutants exhibited significantly reduced fluorescent signals at the junctions (Figure 5D and 5E), despite all three mutants being stably expressed by Western blot analysis (Figure 5F). Because the E291A mutation appeared to be a strong suppressor that is stably expressed and correctly localized, we hypothesized that E291A could be disrupting dimerization of the complex and favoring a monomeric state.

**Figure 5:**
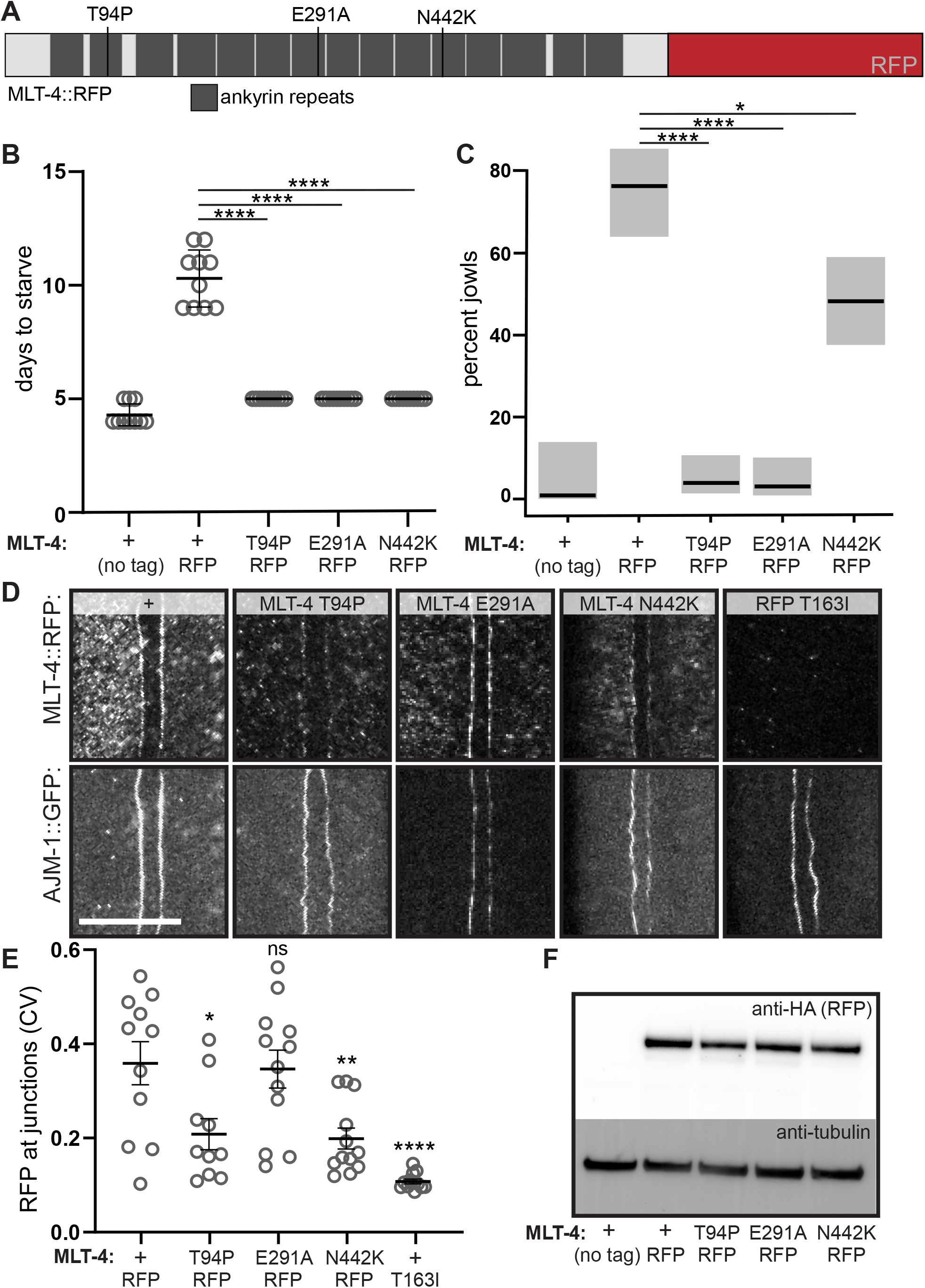
Characterization of missense mutations in MLT-4 that suppress jowls. (A) Schematic of MLT-4::RFP indicating mutations in MLT-4 isolated from mutagenesis screen. (B) Jowls assay. Percent of worms exhibiting jowls; gray boxes represent 95% confidence intervals. n=51-88. (C) Fitness assay. Number of days for population to expand and consume food source. Data represent mean +/-standard deviation of 10 biological replicates. (D) Imaging assay for MLT-4 localization at epithelial junctions. Representative images of endogenously tagged MLT-4::RFP (top) and apical junction marker AJM-1::GFP (bottom). Scale bar = 5 μm. Quantification of MLT-4::RFP pixel intensity at junctions plotted as coefficient of variance (CV). Data represent mean +/-SEM of 10-12 biological replicates. (F) Western blot dual-labeled for HA tag on MLT-4::RFP (top) and loading control (bottom). + = wild-type at *mlt-4* locus. ns p>0.05, * p≤0.05, ** p<0.01, *** p < 0.001, **** p<0.0001, ANOVA analysis with Tukey’s post hoc test, as indicated (B,C) or compared to MLT-4::RFP (E).

If dimerization of the Inversin complex is required for its function, a monomerizing mutant should phenocopy a lethal null mutant in the absence of a dimerization tag (Figure 6A). To test this, we generated the E291A mutant in an untagged MLT-4 strain and compared the phenotype to that of a MLT-4 deletion. Since *mlt-4* nulls are lethal, we generated the MLT-4 deletion and the E291A mutation in worms rescued by extrachromosomal arrays of MLT-4. We then evaluated the phenotype of offspring that did not inherit the array. For both the homozygous MLT-4 null and E291A strains, 0% of the array-negative worms survived, compared to ∼95% of their array-positive siblings (Figure 6B).

**Figure 6:**
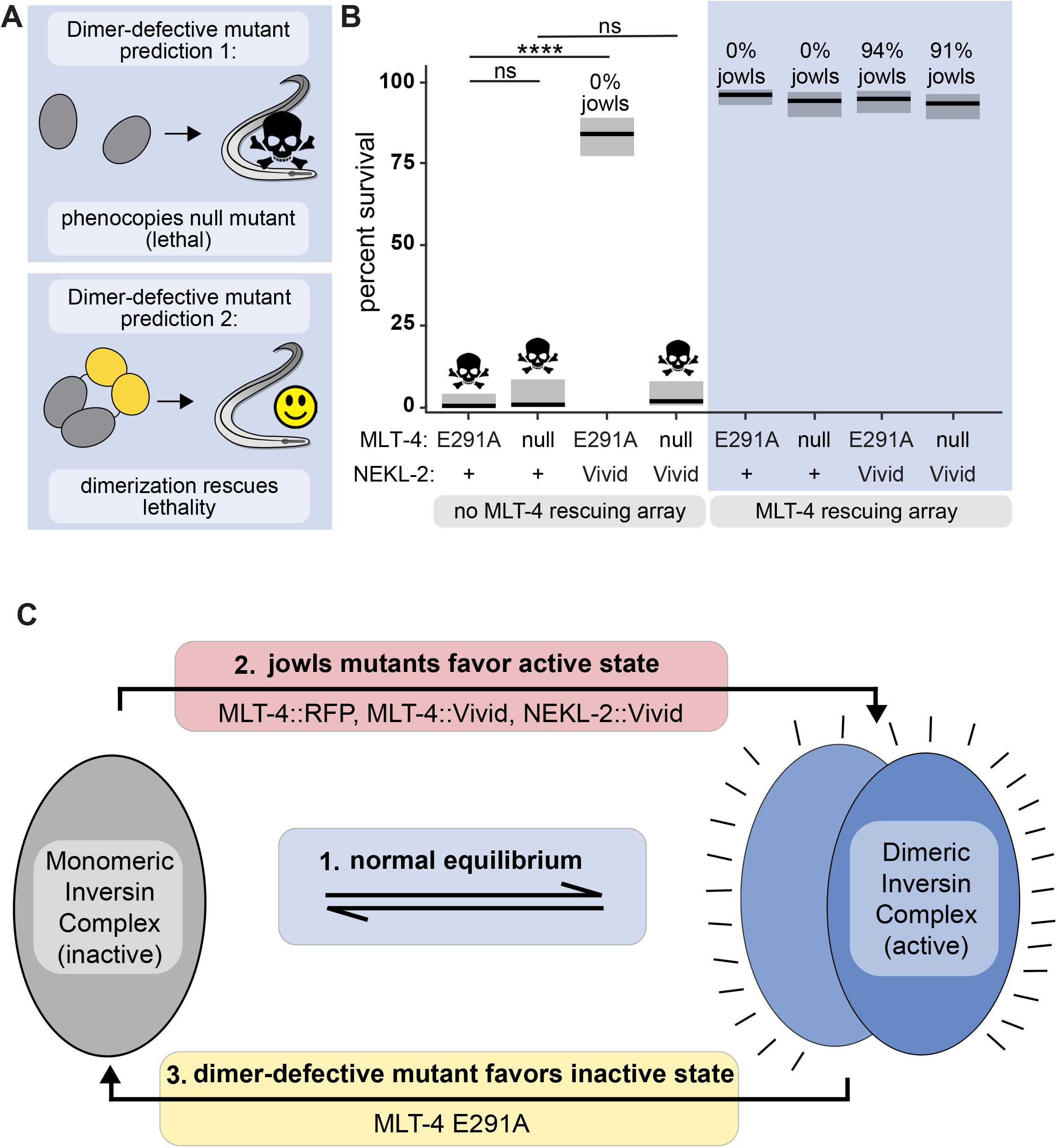
Dimerization of NEKL-2 rescues a functionally inactive MLT-4 mutant. (A) Graphic summary of phenotype predictions for a dimer-defective Inversin complex mutation. (B) Percent of worms surviving to adulthood; gray boxes represent 95% confidence intervals. n=89-271. Percent jowls is indicated for the strains that were viable. (C) Model for dimerization-dependent activation of the Inversin complex. + = wild-type at *nekl-2* locus. ns p>0.05, ** p<0.01, *** p < 0.001, **** p<0.0001, ANOVA analysis with Tukey’s post hoc test, as indicated.

We further reasoned that a monomerizing mutant should be rescued by inducing dimerization of the complex (Figure 6A), whereas a null would not. Presumably, dimerization of MLT-4 by the RFP tag enabled us to isolate the E291A mutation from our screen. To test if dimerization of another member of the complex could similarly rescue the lethality of this mutant, we generated the E291A mutation in the NEKL-2::Vivid strain. We found that light-induced dimerization of NEKL-2 rescued E291A lethality (84% of offspring survived, Figure 6B). Importantly, a MLT-4 deletion was not rescued by dimerization of NEKL-2 (0% of offspring survived, Figure 6B), suggesting that the E291A mutation specifically counteracts dimerization of the Inversin complex.

## Discussion

Loss-of-function analyses and colocalization studies of Inversin complex proteins from both vertebrates and invertebrates show that INVS, NEK8, and ANKS6 work together, but how the activity of the Inversin complex is regulated to produce a functional output is largely unknown. Our results suggest that there are at least two functionally distinct states of the complex: an inactive monomer and an active dimer (Figure 6C). We hypothesize that the hyperactive jowls alleles stabilize the active dimer state; we can induce dimerization optogenetically and recapitulate the phenotype. We further hypothesize that a missense mutation isolated from our suppressor screen appears to favor the inactive monomer state; the lethality of the mutation is bypassed by forcing dimerization of NEKL-2. Thus, we propose dynamic switching between these two states as a new model for how the Inversin complex functions.

### A new functional assay for Inversin complex activity

Previous work has shown that loss of the Inversin complex in *C. elegans* results in a lethal molting defect (Lažetić and Fay 2017; Yochem et al. 2015). In vertebrates, many of the nephronophthisis disease-associated mutations in INVS, ANKS6, and NEK8 result in early stops or frameshifts, suggesting that they also result in loss-of-function (Otto et al. 2003; Kulkarni et al. 2020; Frank et al. 2013). Despite characterization of how these loss-of-function alleles affect localization, ciliogenesis, and cyst formation (Zalli, Bayliss, and Fry 2012; Otto et al. 2008), we lack a comprehensive understanding of the normal activity of the complex due to the absence of gain-of-function analyses, which could identify potential downstream targets. In this paper, we conclude that the MLT-4::RFP allele is likely hyperactive since it is dominant and modulated by the dosage of the wild-type gene (Figure 1D and 1E). This allele enabled us to characterize a recessive point mutation in the NEK8 kinase domain (R12I) associated with end-stage renal failure (Hassan et al. 2020). In our animals, the R12I disease allele suppressed the gain-of-function jowls phenotype, but is not lethal (Figure 1A-1C). These results are consistent with the R12I mutation being a hypomorphic allele, as opposed to a loss-of-function allele. Thus, our jowls assay can be used to assess additional disease-associated missense mutations as they are discovered.

Our results show that constitutive dimerization of MLT-4 or NEKL-2 causes jowls. Although we looked specifically at dimerization in this paper, we cannot rule out that a higher order oligomer is the true active state. Indeed, superresolution microscopy data suggest that the Inversin complex forms fibril-like structures in cilia (Bennett et al. 2020), but the functional significance of these structures remains unclear. Interestingly, we did not observe jowls when we attempted to dimerize MLT-2 by fusing Vivid to either terminus (Figure 4). Several studies in both *C. elegans* and vertebrates suggest that MLT-2/ANKS6 is part of the same complex as NEKL-2/NEK8 and MLT-4/INVS, and is required for activation of the kinase (Hoff et al. 2013; Lažetić et al. 2018; Czarnecki et al. 2015). The simplest explanation for our result is that steric hindrance prevents dimerization of MLT-2 when fused to Vivid. Alternatively, MLT-2 could be functioning differently from MLT-4 and NEKL-2 in our animals to regulate the complex through a mechanism independent of dimerization. Differential impact of dimerization on the individual subunits could also point to hierarchical assembly of Inversin complex components that can be explored in future studies.

Why does the Inversin complex require dimerization to be active? NEK8 is thought to act downstream of INVS (Fukui et al. 2012) and many kinases are known to become active via autophosphorylation in response to dimerization (Hubbard and Miller 2007). A previous study suggests that NEK8 activity is dependent on autophosphorylation (Zalli, Bayliss, and Fry 2012). Our results support a model whereby dimerization could promote autophosphorylation. However, if the kinase were strictly downstream of INVS, then dimerization of the kinase with Vivid should bypass a MLT-4 deletion — it does not (Figure 6B). Instead, this result is consistent with data from multiple systems showing that MLT-4/INVS is required for localization of NEKL-2/NEK8 (Bennett et al. 2020; Lažetić and Fay 2017; Hoff et al. 2013). Indeed, we find that a properly localized, yet lethal, MLT-4 mutant (E291A) is rescued by dimerization of NEKL-2 (Figure 6B). However, the animals did not exhibit jowls, indicating that the E291A mutant reduces dimerization of the kinase. Altogether, these results suggest that the Inversin complex proteins are acting as interdependent subunits.

### An *in vivo* strategy to assess the monomeric character of fluorescent proteins and other genetically encodable tags

Most genetically encodable fluorophores descend from dimeric or tetrameric proteins. Thus, a central focus of fluorescent protein (FP) engineering is generating monomeric versions. The standard *in vitro* assays for determining the oligomeric state of FPs include gel filtration, ultracentrifugation, and structural analyses. However, the propensity of a FP to dimerize often depends on its cellular environment. Currently, the standard *in vivo* assay of FP oligomerization is the OSER assay, which assesses FPs fused to ER-membrane proteins. If the FP oligomerizes, it will deform the ER and create whorl-like structures (Snapp et al. 2003). Using this assay, TagRFP has been shown to dimerize in cells (Costantini et al. 2012), even though it appears monomeric *in vitro* (Merzlyak et al. 2007). Our results are consistent with tagRFP-T having a propensity to dimerize *in vivo* (Figures 2, 3, and 4). We previously found that tagging MLT-4 with mScarlet does not cause jowls (Beacham et al. 2022), which is also consistent with results from the OSER assay where mScarlet exhibited 75% normal cells (Bindels et al. 2017).

Interestingly, our suppressor screen generated mutations in tagRFP-T that have previously been shown to monomerize FPs. Two of the residues mutated in our screen that retain red fluorescence (E205 and V221, Figure 2A) have been previously shown to be important for monomerization of GFP according to both *in vitro* methods and the OSER assay (Zacharias et al. 2002; Costantini et al. 2012; Pédelacq et al. 2006; Scott et al. 2018). The R162 residue in tagRFP-T arose in tagRFP from random mutagenesis, and then later was mutated to reduce dimerization in the development of mKelly2 (Wannier et al. 2018) and fusionRed (Shemiakina et al. 2012). Several other residues that were identified in our suppressor screen as non-fluorescent mutants have also been modified to monomerize other FPs. For example, T110 was mutated to a proline in our screen, but to a lysine in order to generate mCarmine (Fabritius et al. 2018). Similarly, V200E was isolated in our screen but was mutated to isoleucine in the development of mMaroon (Bajar et al. 2016). Our results suggest a novel strategy to evaluate the monomeric nature of any fluorescent protein or other genetically encodable tags *in vivo*.

## Materials and Methods

### Worm strains, maintenance, and CRISPR-Cas9 transgenics

Strains were maintained at room temperature (or 15 °C for Figure 2A) on nematode growth medium (NGM) plates seeded with bacterial food source (strain OP50). CRISPR-Cas9 edits were generated by injecting *C. elegans* gonads with ribonucleoprotein (RNP) complexes as described in (Ghanta and Mello 2020), using unmodified oligonucleotides and slight modifications to the annealing strategy as described in (Beacham et al. 2022). For a complete list of components of the RNP complexes including crRNAs, accompanying repair strategies, and resulting strains and alleles, see Supplementary File 1. Alleles generated by CRISPR were confirmed by sequencing a PCR amplicon of the modified locus.

### MLT-4::RFP suppressor screen

MLT-4::RFP mutants (GUN405) were mutagenized as described in (Beacham et al. 2022). Briefly, worms were incubated for 4 hours at 22 °C in 0.5 mM N-nitroso-N-ethylurea (ENU, Sigma Aldrich N3385), washed thoroughly in M9 buffer (22 mM KH_2_HPO_4_, 42.3 mM Na_2_HPO_4_, 85.6 mM NaCl, 1 mM MgSO_4_), distributed across NGM growth plates seeded with concentrated OP50 bacterial culture, and passaged several times to select for worms with increased fitness. One suppressed animal per culture plate was selected to ensure isolation of independent suppressors. Genomic regions corresponding to the RFP tag and *mlt-4* gene were amplified and sequenced to identify mutations.

### DIC and live fluorescent imaging

For DIC imaging, worms were mounted on 2% agarose pads on glass slides in 20% sodium azide and imaged using a 40x DIC objective 10-20 minutes after overlay of a No. 1.5 glass coverslip. For fluorescent imaging, adult worms were mounted on 8-10% agarose pads on glass slides and immobilized using 0.1 μm polystyrene beads (Polysciences, Warrington, PA, 2.5% by volume) diluted 2-fold in PBS (pH 7.4) followed by overlay of a No. 1.5 glass coverslip (Kim et al. 2013). For initial screening of fluorescent signal in the RFP tag mutants (Figure 2A), the focal planes between the alae and gonad were identified using brightfield microscopy. Then, a Z-stack of the red fluorescent channel was collected using a 60x oil immersion objective on a BZ-X810 Keyence microscope. Worms exhibiting fluorescent signals at epidermal junctions were scored for each strain. For quantification of junctional fluorescent signal (Figures 2, 3, and 5), Z-stacks flanking the AJM-1::GFP junctional marker were collected in both the green and red channels using a custom-built RM21 TIRF microscope (MadCity Labs) equipped with a 60x oil immersion TIRF objective (Nikon), 488 and 552 nm lasers (Coherent OBIS), and an Orca Fusion BT sCMOS camera (Hamamatsu). For image analysis, a segmented line (5 pixels wide) was drawn along the AJM-1::GFP signal and transformed into the red channel using Fiji (Schindelin et al. 2012). The mean pixel intensity and standard deviation (SD) were measured to calculate coefficient of variance (CV = SD / mean) for each segmented line.

### Jowls assays

Synchronized populations of adult worms were scored for the presence of jowls under a dissecting microscope (Nikon SMZ800N). For allele classification in Figure 1E, MLT-4::RFP/– was generated by CRISPR as indicated in Supplementary File 1, resulting in a balanced heterozygous strain (GUN2180). MLT-4::RFP/+ heterozygous worms were generated by crossing N2 males with GUN350 hermaphrodites and selecting F1 animals. Offspring of MLT-4::RFP/+ and MLT-4::RFP/– heterozygotes were scored for jowls and genotyped *post hoc*. Only the confirmed heterozygous offspring were included in the analysis. For phenotypic analysis following induction of RFP::NEKL-2 expression in Figure Supplement 1, jowls were scored 6 hours post heatshock (34 °C, 1 hour).

### Starvation assays

Three L4 worms were placed on each of ten culture plates per strain, and allowed to reproduce and expand. Plates were evaluated daily, and marked as starved when all food was consumed.

### Statistical analysis

Jowls assays and the survival assay in Figure 6 were analyzed in RStudio as described previously (Beacham et al. 2022). Briefly, generalized linear models were fit to a binomial distribution. Starvation assays and imaging assays were analyzed in GraphPad. For all statistical comparisons, ANOVA analysis was performed using Tukey’s method to adjust for multiple comparisons.

### Western blot analysis

For each sample, one-hundred L4 worms were lysed in PBS with 1x Bolt LDS Sample Buffer (Invitrogen) containing 0.01% Triton X-100 and 25 mM DTT. Samples were frozen in liquid nitrogen before sonicating at 70% power for 3 minutes total (1 second on, 1 second off) in a cup horn sonicator. Samples were then incubated at 70 °C for 10 minutes. Sonication and heating steps were repeated 2-3 times to ensure complete lysis. SDS-PAGE was performed using Bolt 4-12% Bis-Tris Plus precast gels (Invitrogen). Proteins were transferred to PVDF Immobilon-FL membranes (Merck Millipore) using the Power Blotter Semi-dry Transfer System (Thermo Scientific) according to product instructions. After transferring, membranes were blocked using EveryBlot Blocking Buffer (Biorad) for 10 minutes. Membranes were incubated in primary antibody diluted in blocking buffer [rat anti-HA HRP 1:500 (Roche 12013819001), mouse anti-tubulin 1:1000 (Sigma, T5168)] for 1 hour at room temperature with orbital shaking, washed 3 times in TBST for 5 minutes each, and incubated in secondary antibody diluted in blocking buffer [goat anti-mouse 488 1:4000 (ThermoFisher Scientific A11029)] for 30 minutes at room temperature with orbital shaking. Before imaging, membranes were washed 3-4 times in TBST for at least 5 minutes each. HRP was detected using SuperSignal West Dura Extended Duration Substrate (Thermo Scientific). Blots were imaged using the BioRad ChemiDoc MP imaging system and band intensities were quantified using ImageLab software.

### Molecular visualization

The structural representation of the tagRFP-T dimer (chains A and B of PDB 5JVA) in Figure 2F was prepared in ChimeraX (Pettersen et al. 2021; Goddard et al. 2018).

### TIRF single molecule imaging

#### Coverslip PEGylation

Glass coverslips were placed in a glass Coplin jar containing acetone and sonicated using a water bath sonicator for 10 minutes. Coverslips were then rinsed 5 times with reverse osmosis (RO) water, sonicated in methanol for 10 minutes, washed 5 times with RO water, sonicated in 3N KOH for 40 minutes, washed 5 times with RO water, and rinsed in methanol. After drying, the coverslips were PEGylated by incubating in a 1:100 mixture of 1% biotin-PEG-silane in ethanol (Laysan Bio Biotin-PEG-SIL-2K-1g) and PEG-silane (85%, VWR 77035-498) for one hour in the dark at room temperature. Coverslips were rinsed thoroughly in RO water and dried with nitrogen gas before storing in the dark at room temperature in a Tupperware container with Drierite desiccant (VWR).

#### *C*. *elegans* lysate preparation

Worms were cultured at room temperature on 15 cm NGM plates seeded with concentrated OP50 *E. coli* culture mixed with chicken egg. When plates were confluent but not yet starved, worms were harvested using TBS and centrifuged at 180 x g at 4 °C for 2 min. Supernatant was removed and the pelleted worms were washed 2 times with TBS. Worms were resuspended in an equal volume of 2x lysis buffer (2x TBS, 0.2% TX-100, and 10% glycerol with protease inhibitors (Roche, 1 tablet per 25 ml lysis buffer)). The resulting worm slurry was frozen dropwise in liquid nitrogen and stored at −80 °C until lysis. Frozen worm pellets were ground in a coffee grinder pre-chilled with liquid nitrogen. When the ground sample lacked intact worms as verified by thawing an aliquot on a slide under a dissecting scope, the remaining powdered sample was stored at -80 °C. Immediately before slide preparation for imaging, an aliquot was thawed on ice and spun at 17,000 × g for 2 min at room temperature. The supernatant was collected and applied to PEGylated coverslips for imaging (described below).

#### Slide preparation for imaging

Prior to imaging, multi-channel devices were assembled by adhering a coated coverslip to an uncoated glass slide using ∼4 mm strips of double-sided tape placed orthogonally to the long axis. Each channel was filled by capillary action with wash buffer (10 mM Tris, pH 8.0, 50 mM NaCl, 0.01% BSA, 0.01% TX-100). Channels were washed 2 times before each of the following steps: 10 minute incubation with 0.2 mg/mL neutravidin (Thermo Scientific) in wash buffer, 10 minute incubation with biotinylated antibody [anti-HA biotin (Roche 12158167001) diluted 1:100 in wash buffer], 10 minute incubation with blocking buffer (10 mM Tris, pH 8.0, 50 mM NaCl, 2% BSA, 0.01% TX-100), and 15 minute incubation with *C. elegans* lysate. Channels were washed 3 times before imaging. Samples were imaged using a RM21 TIRF microscope (MadCity Labs) equipped with a 60x oil immersion TIRF objective (Nikon), a 552 nm laser (Coherent OBIS), and an Orca Fusion BT sCMOS camera (Hamamatsu).

#### Data analysis

Single-molecule detection and analysis of photobleaching steps was performed automatically using SimPull Analysis Software https://github.com/dickinson-lab/SiMPull-Analysis-Software (Stolpner and Dickinson 2022; Dickinson et al. 2017). Samples were excluded if particle densities precluded accurate single molecule spot detection. Single molecules that were rejected by the SimPull Analysis Software (molecules that did not photobleach or molecules that exhibited an increase in intensity over the course of imaging) were excluded from the final analysis.

### Vivid transgenes and light exposure

The sequence of *Neurospora crassa* Vivid (Uniprot entry Q1K5Y8_NEUCR) was synthesized as a gBlock (IDT) with the following modifications: the first 36 amino acids were removed (Zoltowski and Crane 2008; Vaidya et al. 2011), a point mutation (I52C) that has previously been shown to stabilize the dimer state was introduced (Nihongaki et al. 2014), and a *C. elegans* intron was added using *C. elegans* Codon Adapter (Redemann et al. 2011). The resulting synthetic gene (gbEB3) was amplified for CRISPR repairs. Although Vivid is most sensitive to blue light, culture plates were exposed to ambient light because long-term exposure to blue light induces *C. elegans* embryonic lethality.

### Array rescue experiments

Homozygous MLT-4 null and MLT-4 E291A worm strains rescued by MLT-4 extrachromosomal arrays (GUN2409 and GUN2407, respectively) were generated by injecting a MLT-4::GFP plasmid (along with neuronal marker pGH5) into MLT-4::RFP/MLT-4 null and MLT-4::RFP/MLT-4 E291A heterozygotes. Array-rescued MLT-4 null or E291A homozygous candidates (array-positive offspring lacking jowls) were isolated and confirmed by genotyping. The NEKL-2::Vivid; MLT-4 null strain rescued by a MLT-4 array (GUN2413) was generated by crossing heterozygous NEKL-2::Vivid males with GUN2409. The NEKL-2::Vivid; MLT-4 E291A strain (GUN2411) was generated by CRISPR as indicated in Supplementary File 1. For the survival assay in Figure 6, six hour broods were collected from array-positive adults. Upon hatching, siblings were separated into array-positive and array-negative pools, and scored for viability once the array-positive cohort reached adulthood.

## Acknowledgements

We thank the Cornell statistical consulting unit; Kelly Liu for sharing strains; Eric Drier for advice on classification of alleles; Ed Partlow, Chun Han, and Justin Taraska for advice regarding the RFP mutations; Dan Dickinson, Jeff Lange, and Kevin Cannon for advice on single-molecule experiments; Maria Henriquez, Jenna Benton, Kayleigh Morrison, and Mari Camacho for technical support. We also thank the labs of Maurine Linder, Carrie Adler, Josh Chappie, and Rick Cerione for sharing space and reagents, as well as Rick Baker, Kayleigh Morrison, Justin Tapper, and Carrie Adler for constructive criticism that greatly improved the manuscript. This work was supported by a grant from the National Institutes of Health (R01 GM127548) awarded to Gunther Hollopeter.

---

**Figure Supplement 1.**
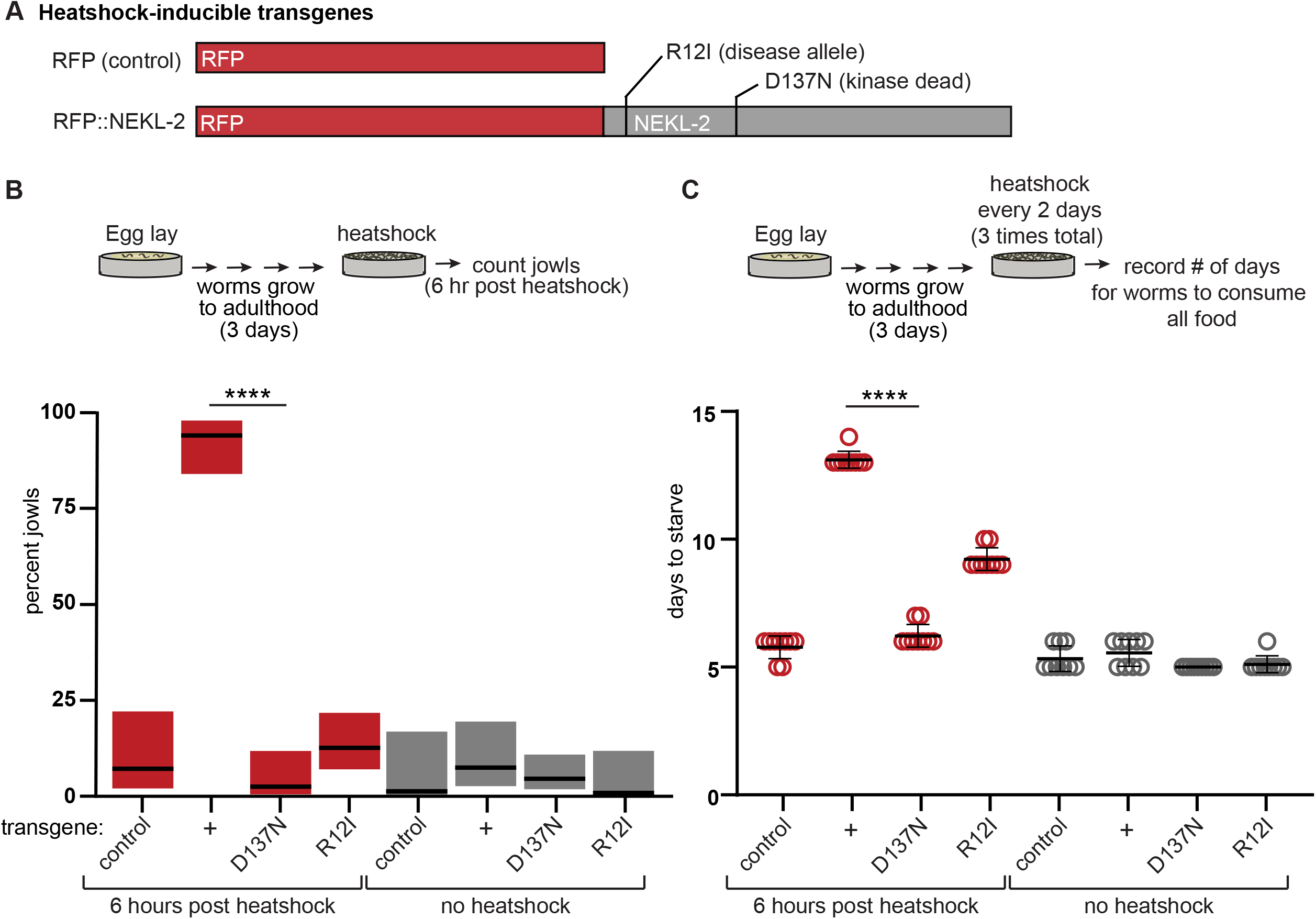
Hyperactive NEKL-2 allele exhibits gain-of-function jowls phenotype. (A) Schematic of heatshock-inducible alleles. (B) Jowls assay. Percent of worms exhibiting jowls; gray boxes represent 95% confidence intervals. n=41-99. (C) Fitness assay. Number of days for population to expand and consume food source. Data represent mean +/-standard deviation of 10 biological replicates. For B and C, control = RFP transgene, + = RFP::NEKL-2 transgene, no mutation. **** p<0.0001, ANOVA analysis with Tukey’s post hoc test, as indicated.

