## Supplemental File 1 for "Dimerization activates the Inversin complex in *C. elegans*"

**Supplementary File 1**

**Strains and alleles**

**Suppressor screen**

GUN405 *ncap-1(mew54[ncap-1::mScarlet::3xflag]) II; mlt-4(mew193[mlt-4::tagRFP-T::HA]) V*

**Figure 1**

Bristol N2 (wild-type)
GUN350 *mlt-4(mew193[mlt-4::tagRFP-T::HA]) V*
GUN775 *nekl-2(mew335[R12I]) I; mlt-4(mew193[mlt-4::tagRFP-T::HA]) V*
GUN652 *mlt-4(mew118[E470K]) V*
GUN706 *nekl-2(mew335[R12I]) I; mlt-4(mew118[E470K]) V*
GUN624 *nekl-2(mew335[R12I]) I*
GUN2180 *mlt-4(mew193[mlt-4::tagRFP-T::HA]) / mlt-4(mew1091[deletion]) V*

**Figure Supplement 1**

GUN565 *mewSi91[Phsp-16.41::HA::tagRFP-T::D10 guide sequence unc-119(+)] IV*
GUN592 *mewSi92[Phsp-16.41::HA::tagRFP-T::nekl-2_cDNA, unc-119(+)] IV*
GUN649 *mewSi93[Phsp-16.41::HA::tagRFP-T::nekl-2(D137N)_cDNA, unc-119(+)] IV*
GUN1006 *mewSi154[Phsp-16.41::HA::tagRFP-T::nekl-2(R12I)_cDNA, unc-119(+)] IV*

**Figure 2**

Imaging strains in 2A:

Mutants from suppressor screen were outcrossed once to remove *mew54[ncap-1::mScarlet::3xflag]*
GUN350 *mlt-4(mew193[mlt-4::tagRFP-T::HA]) V*
GUN442 *mlt-4(mew221[mlt-4::tagRFP-T(I75S)::HA]) V*
GUN443 *mlt-4(mew223[mlt-4::tagRFP-T(L204Q)::HA]) V*
GUN444 *mlt-4(mew225[mlt-4::tagRFP-T(G35R)::HA]) V*
GUN447 *mlt-4(mew231[mlt-4::tagRFP-T(E205K)::HA]) V*
GUN519 *mlt-4(mew233[mlt-4::tagRFP-T(T110P)::HA]) V*
GUN2133 *mlt-4(mew269[mlt-4::tagRFP-T(I10F)::HA]) V*
GUN2134 *mlt-4(mew274[mlt-4::tagRFP-T(R206H)::HA]) V*
GUN2135 *mlt-4(mew273[mlt-4::tagRFP-T(A224V)::HA]) V*
GUN2136 *mlt-4(mew267[mlt-4::tagRFP-T(R162K)::HA]) V*
GUN2137 *mlt-4(mew275[mlt-4::tagRFP-T(V221E)::HA]) V*
GUN2138 *mlt-4(mew281[mlt-4::tagRFP-T(E217D)::HA]) V*
GUN2139 *mlt-4(mew271[mlt-4::tagRFP-T(I10S)::HA]) V*
GUN2140 *mlt-4(mew284[mlt-4::tagRFP-T(T43P)::HA]) V*
GUN2141 *mlt-4(mew288[mlt-4::tagRFP-T(T163I)::HA]) V*
GUN2142 *mlt-4(mew289[mlt-4::tagRFP-T(I10T)::HA]) V*
GUN2143 *mlt-4(mew268[mlt-4::tagRFP-T(V200E)::HA]) V*
GUN2144 *mlt-4(mew285[mlt-4::tagRFP-T(D201G)::HA]) V*

Jowls assay and starvation assay:

Bristol N2 (wild-type)
GUN350 *mlt-4(mew193[mlt-4::tagRFP-T::HA]) V*
GUN1584 *mlt-4(mew707[mlt-4::tagRFP-T(R162K)::HA]) V*
GUN1667 *mlt-4(mew750[mlt-4::tagRFP-T(V221E)::HA]) V*
GUN1709 *mlt-4(mew779[mlt-4::tagRFP-T(E205K)::HA]) V*

Imaging strains in 2D:
GUN731 *ncIs13[ajm-1::gfp]; mlt-4(mew193[mlt-4::tagRFP-T::HA]) V*
GUN2342 *ncIs13[ajm-1::gfp]; mlt-4(mew879[mlt-4::tagRFP-T(T163I)::HA] V*
GUN2271 *ncIs13[ajm-1::gfp]; mlt-4(mew1164[mlt-4::tagRFP-T(E205K)::HA]) V*
GUN2285 *ncIs13[ajm-1::gfp]; mlt-4(mew1179[mlt-4::tagRFP-T(V221E)::HA]) V*
GUN2294 *ncIs13[ajm-1::gfp]; mlt-4(mew1184[mlt-4::tagRFP-T(R162K)::HA]) V*

**Figure 3**
Single-molecule imaging strains:
GUN350 *mlt-4(mew193[mlt-4::tagRFP-T::HA]) V*
GUN1584 *mlt-4(mew707[mlt-4::tagRFP-T(R162K)::HA]) V*

Jowls assay and starvation assay:
Bristol N2 (wild-type)
GUN350 *mlt-4(mew193[mlt-4::tagRFP-T::HA]) V*
GUN1853 *mlt-4(mew876[mlt-4::tagRFP-T(E102A)::HA]) V*

Live imaging strains:
GUN731 *ncIs13[ajm-1::gfp]; mlt-4(mew193[mlt-4::tagRFP-T::HA]) V*
GUN2342 *ncIs13[ajm-1::gfp]; mlt-4(mew879[mlt-4::tagRFP-T(T163I)::HA]) V*
GUN2325 *ncIs13[ajm-1::gfp]; mlt-4(mew876[mlt-4::tagRFP-T(E102A)::HA]) V*

**Figure 4**
Bristol N2
GUN350 *mlt-4(mew193[mlt-4::tagRFP-T::HA]) V*
GUN1321 *mlt-4(mew572[mlt-4::vivid]) V*
GUN1545 *nekl-2(mew688[nekl-2::vivid]) I*
GUN1558 *mlt-2(mew696[mlt-2::vivid]) I*
GUN1748 *mlt-2(mew812[vivid::mlt-2]) I*

**Figure 5**
LW3913 *ncIs13[ajm-1::gfp]*
GUN731 *ncIs13[ajm-1::gfp]; mlt-4(mew193[mlt-4::tagRFP-T::HA]) V*
GUN2342 *ncIs13[ajm-1::gfp]; mlt-4(mew879[mlt-4::tagRFP-T(T163I)::HA]) V*
GUN2235 *ncIs13[ajm-1::gfp]; mlt-4(mew1129[mlt-4(E291A)::tagRFP-T::HA]) V*
GUN2256 *ncIs13[ajm-1::gfp]; mlt-4(mew1150[mlt-4(T94P)::tagRFP-T::HA]) V*
GUN2300 *ncIs13[ajm-1::gfp]; mlt-4(mew1190[mlt-4(N442K)::tagRFP-T::HA]) V*

**Figure 6**
GUN2407 *mewEx166(pKM31, pGH5, Litmus38i); mlt-4(mew1128[E291A]) V*
GUN2409 *mewEx168(pKM31, pGH5, Litmus38i); mlt-4(mew1091[deletion]) V*
GUN2411 *mewEx170(pKM31, pGH5, Litmus38i); nekl-2(mew688[nekl-2::vivid]) I; mlt-4(mew1128[E291A]) V*
GUN2413 *mewEx168(pKM31, pGH5, Litmus38i); nekl-2(mew688[nekl-2::vivid]) I; mlt-4(mew1091[deletion]) V*

| **Allele** | **Generated by** | **Modified locus** | **crRNA** | **CRISPR repair** |
|---|---|---|---|---|
| *mlt-4(mew193[mlt-4::tagRFP-T::HA]) V* | CRISPR (Beacham et al. 2022) | | | |
| *mlt-4(mew118[E470K]) V* | CRISPR (Beacham et al. 2022) | | | |
| *nekl-2(mew335[R12I]) I* | CRISPR | *nekl-2* | rGB246 | oGB582 |
| *mlt-4(mew1091[deletion]) V* | CRISPR | *mlt-4 (mew193)* | rGB325, rGB372 | oEB668 |
| *mewSi91[Phsp-16.41::HA::tagRFP-T::D10 guide sequence unc-119(+)] IV* | CRISPR | *mewSi88[Phsp-16.41::TEV] IV* (Hollopeter et al. 2014) | rEB1, rEB2 | oEB85-oEB86, pEB35 |
| *mewSi92[Phsp-16.41::HA::tagRFP-T::nekl-2_cDNA, unc-119(+)] IV* | CRISPR | *mewSi91* | rEP471 | oEB91-oEB92, pEB36 |
| *mewSi93[Phsp-16.41::HA::tagRFP-T::nekl-2(D137N)_cDNA, unc-119(+)] IV* | CRISPR | *mewSi91* | rEP471 | oEB91-oEB92, pEB39 |
| *mewSi154[Phsp-16.41::HA::tagRFP-T::nekl-2(R12I)_cDNA, unc-119(+)] IV* | CRISPR | *mewSi92* | rGB246 | oGB582 |
| *mlt-4(mew707[mlt-4::tagRFP-T(R162K)::HA]) V* | CRISPR | *mlt-4 (mew193)* | rJZ71 | oEB486 |
| *mlt-4(mew750[mlt-4::tagRFP-T(V221E)::HA]) V* | CRISPR | *mlt-4 (mew193)* | rEB44 | oEB492 |
| *mlt-4(mew779[mlt-4::tagRFP-T(E205K)::HA]) V* | CRISPR | *mlt-4 (mew193)* | rEB42 | oEB537 |
| *mlt-4(mew879[mlt-4::tagRFP-T(T163I)::HA]) V* | CRISPR | *mlt-4 (mew193)* | rJZ71 | oEB587 |
| *mlt-4(mew1164[mlt-4::tagRFP-T(E205K)::HA]) V* | CRISPR | *mlt-4 (mew193)* | rEB42 | oEB537 |
| *mlt-4(mew1179[mlt-4::tagRFP-T(V221E)::HA]) V* | CRISPR | *mlt-4 (mew193)* | rEB44 | oEB492 |
| *mlt-4(mew1184[mlt-4::tagRFP-T(R162K)::HA]) V* | CRISPR | *mlt-4 (mew193)* | rJZ71 | oEB486 |

| | |
|---|---|
| *ncIs13[ajm-1::gfp]* | γ-ray-integrated extrachromosomal array (Z. Liu et al. 2005) |
| *mlt-4(mew221[mlt-4::tagRFP-T(I75S)::HA]) V* | ENU mutagenesis |
| *mlt-4(mew223[mlt-4::tagRFP-T(L204Q)::HA]) V* | ENU mutagenesis |
| *mlt-4(mew225[mlt-4::tagRFP-T(G35R)::HA]) V* | ENU mutagenesis |
| *mlt-4(mew231mlt-4::tagRFP-T(E205K)::HA]) V* | ENU mutagenesis |
| *mlt-4(mew233[mlt-4::tagRFP-T(T110P)::HA]) V* | ENU mutagenesis |
| *mlt-4(mew269[mlt-4::tagRFP-T(I10F)::HA]) V* | ENU mutagenesis |
| *mlt-4(mew274[mlt-4::tagRFP-T(R206H)::HA]) V* | ENU mutagenesis |
| *mlt-4(mew273[mlt-4::tagRFP-T(A224V)::HA]) V* | ENU mutagenesis |
| *mlt-4(mew267[mlt-4::tagRFP-T(R162K)::HA]) V* | ENU mutagenesis |
| *mlt-4(mew275[mlt-4::tagRFP-T(V221E)::HA]) V* | ENU mutagenesis |
| *mlt-4(mew281[mlt-4::tagRFP-T(E217D)::HA]) V* | ENU mutagenesis |
| *mlt-4(mew271[mlt-4::tagRFP-T(I10S)::HA]) V* | ENU mutagenesis |
| *mlt-4(mew284[mlt-4::tagRFP-T(T43P)::HA]) V* | ENU mutagenesis |
| *mlt-4(mew288[mlt-4::tagRFP-T(T163I)::HA]) V* | ENU mutagenesis |
| *mlt-4(mew289[mlt-4::tagRFP-T(I10T)::HA]) V* | ENU mutagenesis |
| *mlt-4(mew268[mlt-4::tagRFP-T(V200E)::HA]) V* | ENU mutagenesis |
| *mlt-4(mew285[mlt-4::tagRFP-T(D201G)::HA]) V* | ENU mutagenesis |

| | | | | |
|---|---|---|---|---|
| *mlt-4(mew876[mlt-4::tagRFP-T(E102A)::HA]) V* | CRISPR | *mlt-4 (mew193)* | rJZ72 | oEB585 |
| *mlt-4(mew572[mlt-4::vivid]) V* | CRISPR | *mlt-4* | rEP739 | oEB326-oEB327, pEB77 |
| *nekl-2(mew688[nekl-2::vivid]) I* | CRISPR | *nekl-2* | rGB423 | oEB459-oEB460, pEB77 |
| *mlt-2(mew696[mlt-2::vivid]) I* | CRISPR | *mlt-2* | rEP1062 | oEB464-oEB394, pEB77 |
| *mlt-2(mew812[vivid::mlt-2]) I* | CRISPR | *mlt-2* | rGB676 | oEB506-oEB507, pEB77 |
| *mlt-4(mew1129[mlt-4(E291A)::tagRFP-T::HA]) V* | CRISPR | *mlt-4 (mew193)* | rGB544 | oGB545 |
| *mlt-4(mew1150[mlt-4(T94P)::tagRFP-T::HA] V* | CRISPR | *mlt-4 (mew193)* | rGB542 | oGB543 |
| *mlt-4(mew1190[mlt-4(N442K)::tagRFP-T::HA]) V* | CRISPR | *mlt-4 (mew193)* | rEB48 | oEB505 |
| *mlt-4(mew1128[E291A]) V* | CRISPR | *mlt-4* | rGB544 | oGB545 |
| *mewEx166(pKM31 [10 ng/µl], pGH5 [20 ng/µl], Litmus38i [70 ng/µl]* | Extrachromosomal array | | | |
| *mewEx168(pKM31 [10 ng/µl], pGH5 [20 ng/µl], Litmus38i [70 ng/µl]* | Extrachromosomal array | | | |
| *mewEx170(pKM31 [10 ng/µl], pGH5 [20 ng/µl], Litmus38i [70 ng/µl]* | Extrachromosomal array | | | |

**CRISPR guides, oligonucleotides, plasmids, and gBlocks**

CRISPR guides:

rEB1 CATCTAAAAAACTTCGAAAA
rEB2 ACTAAATCTGAGCTCCGCAT
rEB42 TTATTACGTGGATCACCGTC
rEB44 CTATGTGGAACAACATGAAG
rEB48 CAACATGAATGACGTAGACG
rEP471 GCTACCATAGGCACCACGAG
rEP739 TTTAAGAAAGAAATAAAAAC
rEP1062 TTACTGAAATCTACAGTGAT
rGB246 AAAGTGCGTGTCGTCGGACG

rGB325 TTTAAATATTGATTTATAAA
rGB372 CGTAGTCAGGGACGTCGTAT
rGB423 AAGAAACAAACAATTAATAC
rGB542 AGTTGTACTGGGTAGCCCAG
rGB544 AATTTCCGGGTAGCCTTGCT
rGB676 ATTTTTTGTAAAGAAATGAG
rJZ71 CCATATCAGTTCTGCCTTCC
rJZ72 CGTGTGACCACTTATGAAGA

Oligonucleotides:
oEB85 ACGCTTTGTTCTAGTGCATCTAAAAAAC
oEB86 GGCGCGAGATGGCGAT
oEB91 CGGCCGATTTAGCTACCATAGGCACCACGTTAATACTTTGAATGCACTTGACTTCTTGAT
oEB92 CCCTAGCAAACTGGGGCACAAACTTAATTCCGCTGGTGGCACAGGAGGAACG
oEB326 CGTTATAAGGAATTTATTTTAAGAAAGAAAAGTACTAGCGG
oEB327 AAAAAAAAAATTAAAATTTTTTTAATAGTTCCTGTTTCTACTCGGTCTCGCATTGGAATC
oEB394 TTAGAATTAATGGCGTTTACTGAAATCTACTCGGTCTCGCATTGGAATCC
oEB459 ATTTTAATTTTAAAGAAACAAACAATTAATTCACGTGTTACTCGGTCTCGCATTGGAATC
oEB460 AAGAAGTCAAGTGCATTCAAAGTATAGTACTAGCGGTGGCAGTGG
oEB464 ATTGAGGATATTCAAACTGCAATTCTTAGCCGATCTCTACACACCCTCTACGCCCCA
oEB486 GCTGTATCCAGCTGATGGTGGCCTCGAGGGTAAAACTGATATGGCACTGAAACTGGTGGG
oEB492 CGATAAAGAAACCTATGTGGAACAGCACGAGGAAGCTGTCGCTCGTTACTGCGATCTGCC
oEB505 GGCCTCCGGAGCCAACATGAAAGACGTCGACGAGGTGGGACGTACGGCGGTCTTCTGGGC
oEB506 TCGATTATTCCCTGAGATTTTTTGTAAAGAAATGCACACCCTCTACGCCCCA
oEB507 CTCTATCGTCAATTCCATCTGCTGCAGCACCGCTCTCGGTCTCGCATTGGAATCC
oEB537 AATGCCGGGAGTTTATTACGTGGATCACCGCCTTAAGCGCATCAAAGAAGCCGATAAAGA
oEB585 TTCACATGGGAACGTGTGACTACGTACGCAGACGGTGGAGTCCTGACAGCAACGCAGGAT
oEB587 GCTGTATCCAGCTGATGGTGGCCTCGAGGGTAGAATTGATATGGCACTGAAACTGGTGGG
oEB668 TTTATCATATTTTTTAAATATTGATTTATAACGACGTCCCTGACTACGCTTAAAAACAGG
oGB543 ATTATAATAAAGCTACCGTACTCCACTGGGCTCCCCAGTACAATTGCAAGCAGGTATCTT
oGB545 CAAAGGACTCCGTTGCACTATGCCGCTGCGCAAGGCTACCCGGAAATTGTGAAATTTCTC
oGB582 GACAATTATGAAAAAGTGCGTGTCGTCGGAATTGGCGCCTTTGGAGTTTGTTGGCTGTGT

Plasmids:
pEB35 Phsp16.41:flexible_linker:HA-tag:flexible_linker:tagRFP-T:D10 guide sequence
pEB36 flexible_linker:NEKL-2 cDNA
pEB39 flexible_linker:NEKL-2 D137N cDNA
pEB77 Ppdi-1:flexible_linker:Vivid
pKM31 Psemo-1:MLT-4 cDNA:flexible_linker:GFP:strep-tag:let-858 3’ UTR
pGH5 Punc-17:GFP:unc-54 3’ UTR
Litmus38i (NEB #N3538)

gbEB3: Vivid amino acid sequence used for CRISPR repairs
CACACCCTCTACGCCCCAGGAGGATACGACATCATGGGATACCTCTGCCAAATCATGAAC
CGTCCAAACCCACAAGTCGAGCTCGGACCAGTCGACACCTCCTGCGCCCTCATCCTCTGC
GACCTCAAGCAAAAGGACACCCCAATCGTCTACGCCTCCGAGGCCTTCCTCTACATGACC
GGATACTCCAACGCCGAGGTCCTCGGACGTAACTGCCGTTTCCTCCAATCCCCAGACGGA
ATGGTCAAGCCAAAGTCCACCCGTAAGTACGTCGACTCCAACACCATCAACACCATGCGTA
AGgtaagtttaaacatatatatactaactaaccctgattatttaaattttcagGCCATCGACCGTAACGCCGAGGTCCA
AGTCGAGGTCGTCAACTTCAAGAAGAACGGACAACGTTTCGTCAACTTCCTCACCATGATC
CCAGTCCGTGACGAGACCGGAGAGTACCGTTACTCCATGGGATTCCAATGCGAGACCGAG
